# Buyang Huanwu Decoction Ameliorates Peripheral Nerve Injury by Promoting Cell Cycle through Activation of PI3K/AKT/mTOR/HIF-1α Pathway

**DOI:** 10.1101/2025.07.02.662709

**Authors:** Honghui Li, Xudong Zhou, Pengtao Huang, Xiaoqing Zhang, Yuping Gao, Bifeng Fu

**Affiliations:** Spine Orthopedics Ward, The First Hospital of Hunan University of Chinese Medicine, Changsha, 410007, Hunan, China; School of Pharmacy, Hunan University of Chinese Medicine, Changsha, 410208, Hunan, China; Department of Traditional Chinese Orthopedics and Traumatology Ward, National Clinical Research Center for Chinese Medicine Acupuncture and Moxibustion, The First teaching Hospital of Tianjin University of Traditional Chinese medicine, 88 Changling Road, Xiqing District, 300381, Tianjian, China; Clinical College of Traditional Chinese Medicine, Tianjin University of Traditional Chinese Medicine, Tianjin, 301617, China

**Keywords:** BHD, Peripheral nerve injury, Cell cycle, p27Kip

## Abstract

**Background:** Peripheral nerve injury (PNI) can lead to partial or even total motor and sensory dysfunction, severely affecting the work and life of patients, but recovery of nerve function is often poor due to the complexity of the nerve repair and regeneration process. Cumulative evidence suggests the contribution of cell cycle regulation in repairing PNI injury, and p27KIP1 is a key molecule in regulating the cell cycle.

**Objective:** The aim of this study is to elucidate the repair mechanism of peripheral nerve injury (PNI) from the point of view of regulating cell cycle and to explore the therapeutic effect of Buyang Huanwu Decoction (BHD). To provide new strategies for the treatment of PNI and to explore valuable drugs initially.

**Methods:** The clamp injury model was chose as a PNI model and different doses of BHD extract were administered by gavage for 14 days. Limb motor function, sciatic nerve index and sciatic nerve pathology were assessed and p27Kip expression level was determined. The in vivo chemical composition of BHD extract was determined and drug targets were retrieved by TCMSP, STITCH and Swiss Target Prediction to predict the mechanism of BHD treatment for PNI using network pharmacology. Pathway-related proteins including P-PI3K, P-AKT, P-mTOR and HIF-1a were validated by western blotting assay.

**Results:** Compared with control rats, PNI rats had abnormal motor function, sciatic nerve damage and reduced p27Kip expression. Administration of BHD for 14 days significantly improved PNI in rats, which not only repaired sciatic nerve function, attenuated sciatic nerve pathology and inhibited p27Kip expression to regulate the cell cycle in PNI rats, but also activated the PI3K/AKT/mTOR/HIF-1a pathway.

**Conclusion:** The above results suggest that PI3K/AKT/mTOR/HIF-1a mediated cell cycle regulation may be an important repair mechanism for PNI, and that BHD can improve PNI by accelerating the cell cycle to promote neural repair and regeneration.

## Introduction

Peripheral nerve injury (PNI) is the injury of nerves outside the brain and spinal cord usually caused by war, disease, traffic accidents^[1, 2]^. Peripheral nerve injury can lead to partial or even total motor and sensory dysfunction, resulting in varying degrees of lesions in the peripheral nervous system and the central nervous system, which can seriously affect the patient’s work and life and cause a serious financial burden on the family ^[1–5]^. Therefore the treatment of PNI has been of great interest to researchers. Currently, treatments for PNI include nerve growth factors, stem cell-derived exosomes, physical therapy, nerve conduit or scaffolding materials, and drug therapy ^[1, 2, 6]^. However, due to the complexity of the process of nerve repair and regeneration and the long time required, the process due to a variety of factors, the recovery of nerve function is often not good ^[7–9]^, so the search for PNI intervention method points and therapeutic drugs is so medical workers need to pay attention to and solve the problem.

Cell cycle regulation refers to the regulation of cell cycle progression through the interaction of cell cycle protein-dependent kinase (CDK) and cell cycle protein-dependent kinase inhibitor (CDKI), which plays an important role in the process of neural regeneration and repair ^[10–12]^. Schwann cells (SCs), as glial cells unique to the peripheral nervous system, play a key role in the repair and treatment of PNI ^[13]^. Under normal conditions, they maintain the functionality of the nerve by regulating the axon diameter, the distribution of ion channels and the phosphorylation of nerve microfilaments, and after the peripheral nerve injury, the SCs are transformed into a reparative phenotype and proliferate in a large number of divisions and secrete cytokines as well as neurotrophic factors, providing a microenvironment conducive to nerve regeneration, providing nutrient support and physical protection for the regenerated axons^[14, 15]^. p27KIP1 is a key molecule in the regulation of the cell cycle and is involved in the regulation of neuronal cell proliferation. Wang et al. found reduced p27kip1 expression in both neurons and glial cells in the PNI state, and hypothesized that the reduced p27kip1 levels were associated with axonal regeneration and glial cell proliferation ^[12]^.Cumulative studies have also reported that p27KIP1 inhibition of Cdk-cyclin kinase and CDK activity is attenuated in the PNI state, and increases the activity of transcription factors and accelerates the process of the cell cycle and thus participates in the regeneration and repair of nerves in the PNI state^[16–20]^. In addition, it was found that p27 expression levels were reduced in a TNF-α-induced Schwann cell proliferation model, demonstrating that P27KIP is involved in the proliferation and differentiation of SCs after sciatic nerve crush ^[21]^. Thus, p27Kip-1, as a protein involved in cell cycle regulation, plays an important role in the proliferation of SCs and the repair of PNI.

According to PNI clinical manifestations, Chinese medicine often categorizes it as “impotence” and “paralysis”, and believes that its pathogenesis is mainly due to the blockage of meridians and channels, deficiency of qi and blood stasis. Buyang Huanwu Decoction (BHD) is consists of seven herb including Astragali Radix (*Astragalus membranaceus* ( Fisch.) Bge., huangqi), Angelica Sinensis Radix (*Angelica sinensis* (Oliv.) Diels, danggui), Paeoniae Radix (*Paeonia lactiflora* Pall., chishao), Carthami Flos *(Carthamus tinctorius* L., honghua), Persicae Semen (*Prunus persica* (L.) Batsch, taoren), Chuanxiong Rhizoma (*Ligusticum chuanxiong Hort.*, chuanxiong) and Pheretima (*Pheretima aspergillum* (E.Perrier), dilong). BHD effects of tonifying qi, invigorating blood and clearing collaterals have a good therapeutic effect on PNI. Studies on the clinical use of BHD in the treatment of PNI have also been reported, and it has been found to be effective and safe during treatment, and can be a complementary and alternative option to monotherapy with neurotrophic drugs or electrical stimulation^[22]^, however, there is a lack of research on the mechanism of action by which it exerts its efficacy that needs to be further elucidated. Therefore, this study attempted to explore the effect of BDH on PNI from the perspective of regulating cell cycle to promote nerve regeneration to provide new strategic ideas and approaches for clinical treatment.

## Materials and methods

### Preparation of BHD extract

BHD consists of Astragali Radix 120g, Angelica Sinensis Radix 6g, Paeoniae Radix 5g, Chuanxiong Rhizoma 3g, Persicae Semen 3g, Carthami Flos 3g, and Pheretima 3g. All herbal medicines were purchased from the First Affiliated Hospital of Hunan University of Traditional Chinese Medicine. All the herbs were washed through pure water, added 10 times distilled water and soaked for 30 min, boiled for 0.5 h on military fire, slow decoction for 1 h on civil fire. The filtrate was collected, and the residue was extracted again with 5 times the amount of distilled water in the same way for 3 times. The filtrates were combined, concentrated to 2 g/ml, and stored in a -80° refrigerator.

### Animals and Experimental design

40 SD male rats were purchased from Hunan SJA Laboratory Animal Co., Ltd (SCXK(Xiang)2019-0004), weighing 220g∼240g. All experiments were approved by the Ethics Committee of the First Hospital of Hunan University of Traditional Chinese Medicine(ZYFY20210507-11), and all operations complied with the ethical requirements for laboratory animals. The rats were deeply anesthetized using isoflurane after 3 days of acclimatization feeding, and after the rats were successfully anesthetized by intraperitoneal injection, the surgical area was disinfected and treated in the right buttock of the rats. Surgical operations in the model group and the drug administration group were as follows: a 2 cm incision was made obliquely in the right buttock of the rats, and the skin, subcutaneous tissue and deep fascia were incised layer by layer to bluntly detach the gluteal muscles of the rats, and the sciatic nerve on the right side was fully exposed. The sciatic nerve was clamped using a microscopic hemostat 5 mm from the inferior border of the pyriformis muscle. Clamping was required to the extent of a second buckle, maintained for 10 s each time, and repeated 3 times at 5-s intervals. The rat sciatic nerve injury model was prepared by ensuring that the pressure reached 21,950 Pa, resulting in an area of nerve injury approximately 2 mm wide. In the sham-operated group, no clamping operation was performed, and the rest of the steps were consistent. After the experimental operation, rats were divided into control group, PNI group, BHD low dose group (BHDL, 7.15g/kg), BHD high dose group (BHDH, 14.3g/kg) and positive drug group(methylcobalamin, 0.15 mg/kg), the treatment group was gavaged with the corresponding dose of BHD extract twice a day for 14 days.The rats in Con group and PNI group were given equal dose of distilled water by gavage. Positive drug group was administered with positive drug methylcobalamin (Lot No. 1908028, Eisai (China) Pharmaceutical Co.).

### Measurement of limb motor function

To test the effect of BHD on limb motor function in rats with PNI, sciatic nerve recovery was measured on postoperative days 3, 7, and 14 using the slant plate test and sciatic nerve function index (SFI). The inclined plate experiment was conducted by raising the plank from 0°for every 5°, if the rat persisted for 5s continue to raise the plank for 5° until the rat could not persist then the current data was recorded. Each rat rested for 3-5 min after measurement and repeated the experiment again, for a total of three times. The sciatic function index (SFI) Select the footprints of the experimental side foot (E) and the normal side foot (N), select 3 or 4 footprints on each side, and measure 3 variables respectively: (1) Print length (PL): the distance from the heel to the tip of the foot. (2) Distance between first and fifth toes or total spreading (TS). (3) Distance between intermediate toes (IT) from the 2nd toe to the 4th toe. SFI=-38.3 [(Ep L-Np L)/Np L]+109.5 [(ETS NTS)/NTS]+13.3 [(EIT NIT)/NIT] -8.8. Bring the above three variables into the Bain formula to calculate SFI^[23]^. The sciatic nerve function index (SFI)=0 indicates normal, and -100 indicates complete injury.

### Hematoxylin and eosin

To evaluate the effect of BHD on the recovery of sciatic nerve, after execution of rats under deep anesthesia, the sciatic nerve on the operated side of rats was taken, fixed with 4% paraformaldehyde, dehydrated, and then subjected to 4 μm serial paraffin sections. Experiments were performed with hematoxylin andeosin staining on transverse sections versus longitudinal sections of the sciatic nerve. The stained sections were visualized and captured under a biomicroscope (DM4000B, Leica).

### Immunohistochemistry analysis

To assess the changes of cell cycle-related protein p27kip1 in neural tissues using immunohistochemical staining. Paraffin sections were soaked in 3% H2O2 for 5 min to remove endogenous catalase activity, and non-specific antibodies were blocked with normal goat serum. Diluted primary antibody (Cat. No. ab32044, abcam) was added dropwise and incubated at 4° for 12h, followed by dropwise incubation with appropriate amount of goat anti-rabbit IgG antibody (Cat. No. ab6702, abcam) for 30min at room temperature. Finally, ready-to-use DAB (Cat. No. P0202, beyotime) were added for color development, the staining was terminated after the appearance of positivity, and hematoxylin (Cat. No. C0107, beyotime) were added for restaining. Sections were sealed and visualized and captured under a biomicroscope (DM4000B, Leica).

### Western blot analysis

In order to study the effect of BHD on cell cycle-related proteins in rats with PNI, we used deep anesthesia followed by cervical dislocation to execute the rats, and then collected and took the sciatic nerve on the surgical side of the rats, and stored it at -80°C waiting for detection. Total protein was extracted and quantified with a BCA protein quantification kit according to the manufacturer’s protocols^[23, 24]^. Briefly, 20 mg of sciatic nerve specimen was taken, lysed with 100 ul of lysis solution containing 10 ul of PMSF, centrifuged at 12,000 g for 5 min at 4°C, and the supernatant was collected, and the protein concentration was determined using the BCA Protein Quantification Kit (Cat. No. P0012, beyotime). Total proteins were then separated by polyacrylamide gel electrophoresis (SDS-PAGE) and transferred to a polyvinylidene difluoride (PVDF) membrane.The immunoblotted was then incubated overnight at 4°C with primary antibody: anti-PI3 Kinase(Cat. No.4249S, Cell Signaling Technology) antibody,anti-Phospho-PI3 Kinase(Cat. No.17366S, Cell Signaling Technology) antibody, anti-Akt(Cat. No.9272S, Cell Signaling Technology) antibody, anti-Phospho-Akt(Cat. No.4060S, Cell Signaling Technology) antibody, anti-mTOR Antibody(Cat. No.2972S, Cell Signaling Technology) antibody, anti-Phospho-mTOR Antibody(Cat. No.2971S, Cell Signaling Technology) antibody, anti-HIF-1α Antibody(Cat. No.14179S, Cell Signaling Technology) antibody,anti-p27 Kip1 Antibody(Cat. No.2552S, Cell Signaling Technology) antibody and anti-GAPDH (Cat. No.5174S, Cell Signaling Technology) antibody. After 3 washes, the immunoblots were incubated with peroxidase-conjugated secondary antibodies for 2 hours at room temperature. Proteins were detected using an enhanced chemiluminescence detection kit (Millipore). Band intensities were quantified using Image J.

### UPLC-Q-TOF/MS analysis of compounds in BHD extract

To identify the chemical composition of BHD extracts, UPLC-Q-TOF/MS was used for analysis. The BHD extract at a concentration of 1 g/ml was filtered through a 0.22 μm membrane filter. Mixed standards were prepared at a concentration of 1 mg/ml. The separation was carried out on an ACQUITY UPLC® HSS T3 (2.1×150 mm, 1.8 µm) (Waters, Milford, MA, USA) column at 40 °C with a 2 μL injection volume. In positive ion mode, the mobile phase was 0.1% formic acid acetonitrile (C) and 0.1% formic acid water (D), and the gradient elution program was as follows: 0-1 min, 2%C; 1∼9 min, 2%∼50% C; 9∼12 min, 50%∼98% C; 12∼13.5 min, 98%C; 13.5-14 min, 98%-2% C; 14 to 20 min at 2% C. In negative ion mode, the mobile phase was acetonitrile (A) and 5 mM ammonium formate water (B), and the gradient elution program was as follows: 0-1 min, 2%A; 1-9 min, 2%-50% A; 9-12 min, 50%-98% A; 12∼13.5 min, 98% A; 13.5-14 min, 98%-2%a; 14 to 17 min, 2% A. The MS parameters were as follows: Thermo Q Exactive mass spectrometry detector (Thermo Fisher Scientific, USA), electrospray ion source (ESI), and positive and negative ion modes were used to collect data separately. The positive ion spray voltage was 3.50 kV, the negative ion spray voltage was -2.50 kV, the sheath gas was 30 arb, and the auxiliary gas was 10 arb. The capillary temperature was 325 ℃, the first stage full scan was performed with a resolution of 70000, the first stage ion scanning range was m/z 100-1000, and the second stage lysis was performed with HCD, the collision energy was 30 eV, the second stage resolution was 17500, and the first 10 ions were cleaved before the signal acquisition. At the same time, dynamic exclusion was used to remove unnecessary MS/MS information solution preparation.

### Predicting targets related to PNI

In the OMIM, GeneCards, and DisGeNET databases, gene target queries were conducted using the keyword “Peripheral Nerve Injury” to obtain the currently identified targets for cervical spondylotic radiculopathy, delete duplicate targets, and determine the final target.

### Constructing the drug-compound-disease-target network and identification of core targets

Importing crossover genes into the STRING database, set the protein type to “homo sapiens” ^[25]^, and obtain an information map of protein interactions. Set the score of the lowest confidence protein parameter to “>0.4”, removed edge free proteins, ensured the reliability of the research data, and exported it as a tsv file for backup. The determination of core proteins was based on node degree. The active ingredients and cross genes of BHD were introduced into Cytoscape3.9 to construct a drug-ingredient-disease network, from which the core ingredients for the treatment of PNI were screened.

### GO functional enrichment analysis and KEGG pathway enrichment analysis

GO functional enrichment analysis and KEGG pathway enrichment analysis were performed on the proteins in the “active ingredient potential target” network of the “BHD” drug pair associated with PNI, and significant enrichment of signal pathways was obtained (P<0.05). The above process applies DIVID6.8 ( https://david.ncifcrf.gov/) Software.

### Statistical analysis

Data are expressed as mean ± standard deviation (SD). Statistical analysis was performed using GraphPad Prism 7 software (GraphPad software, Inc., La Jolla, CA, United States), and all data comparisons were performed using one-way analysis of variance (ANOVA). P< 0.05 was considered statistically significant.

## Results

### BHD improves sciatic nerve function in PNI rats

To investigate the effects of herbal compound BHD on limb function in rats with PNI, motor function recovery and SFI were measured on days 3, 7, and 14 days after drug intervention. The results showed that the oblique plate experimental angle was significantly decreased (Fig. 1A) (*P*<0.001) and SFI was significantly decreased (Fig. 1B) (*P*<0.001) in PNI rats compared with the normal group. After treatment with BHD and methylcobalamin, the results showed a significant increase in oblique plate experimental angle in the BHDH (Fig. 1A)(*P*<0.01)and methylcobalamin (Fig. 1A)(*P*<0.05) groups on 3 and 7 days. Significant increase in oblique plate experimental angle in BHDL (Fig. 1A)(*P*<0.05), BHDH (Fig. 1A)(*P*<0.001) and methylcobalamin (Fig. 1A)(*P*<0.05) group on 14 days. And a significant increase in SFI in the BHDL, BHDH and methylcobalamin (Fig. 1B)(*P*<0.001) groups (Fig. 1B)(*P*<0.001) on 3, 7 and 14 days. To further explore the therapeutic effect of BHD on sciatic nerve injury in rats with PNI, pathological changes in sciatic nerve injury (Fig. 1C-D) were detected by HE staining in each group after 14 days of drug intervention, and the results showed that the sciatic nerve of rats in the control group had no obvious pathologic changes, and the myelinated nerve fibers were arranged in an orderly and dense way, with a normal morphology and structural integrity. Structural abnormalities were observed in the sciatic nerve of PNI rats, with disorganized and loosely arranged nerve fibers, more scattered cavities and loss of myelin sheaths, axonal atrophy, and an increase in Schwann cells. The sciatic nerve lesions after BHD and methylcobalamin treatment were improved to different degrees, the nerve fibers were arranged more regularly, the myelin sheath loss and axonal atrophy were significantly improved, and the proliferation and differentiation of Schwann cells, and the therapeutic effect of BHDH was better than that of BHDL. The above results indicated that BHD could significantly improve the sciatic nerve function of rats with PNI.

**Figure 1.**
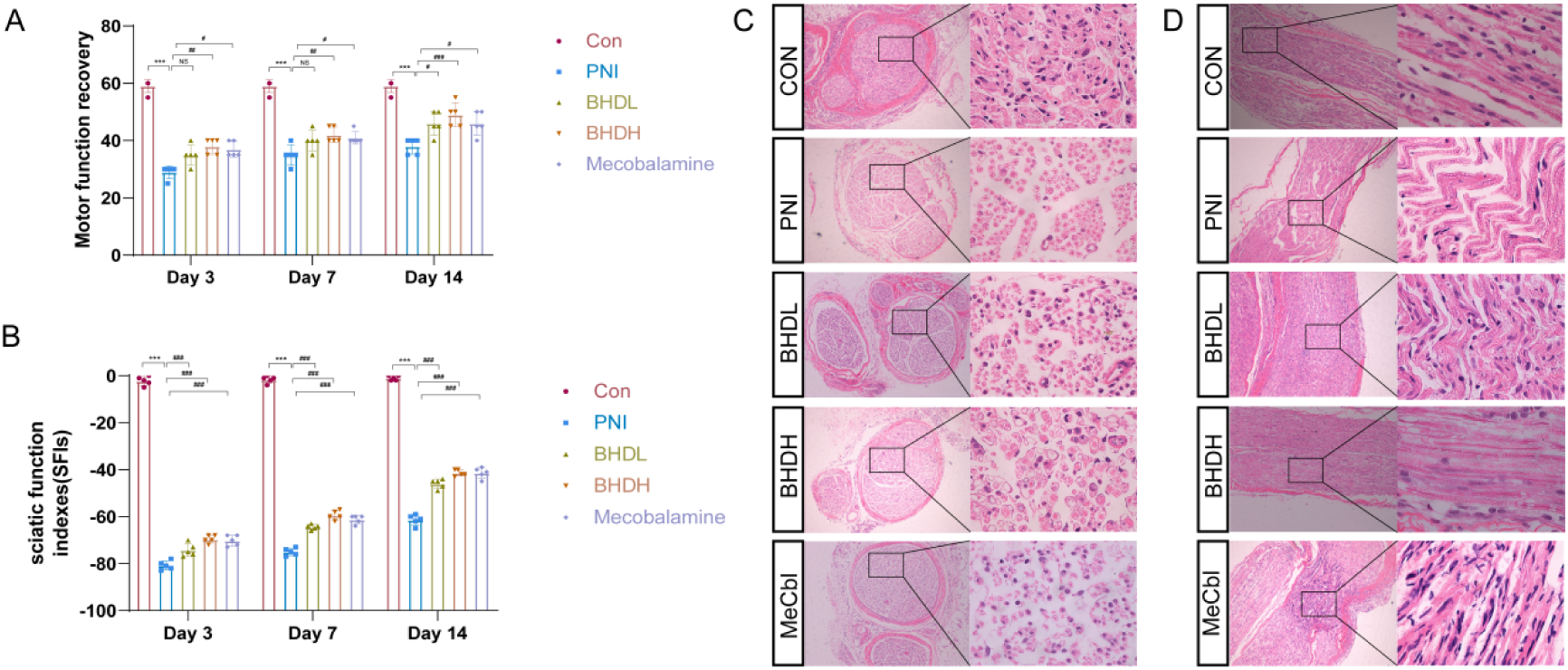
BHD improves sciatic nerve function in PNI rats. (A) motor function recovery, (B) sciatic function index,these data were expressed as the mean ± SD, (n=5 for each group). (C) BHD improved the pathological damage of sciatic nerve cross section in PNI rats (n = 3). (D) BHD improved the pathological injury of longitudinal section of sciatic nerve in PNI rats (n = 3). All data were analyzed by one-way ANOVA, ***P< 0.001,**P< 0.01, *P < 0.05 vs. Con group, ###P < 0.01, ##P < 0.01, #P < 0.05 vs. PNI group.

### BHD decreases p27Kip1 protein expression in sciatic nerve of PNI rats

Given that the proliferation of Schwann cells plays an important role in the treatment of PNI, and in recent years, it has been found that the p27Kip1 protein may promote the proliferation of Scs by regulating cell cycle progression. We examined the sciatic nerve p27Kip1 content in different groups of rats, and the results showed that the sciatic nerve p27Kip1 content in PNI rats was significantly decreased compared to Con group (Fig. 2A-D), while p27Kip1 decreased after BHDL, BHDH, and methylcobalamin treatments compared to the PNI group (Fig. 2A-D). The results showed that p27Kip1 protein expression was increased in PNI rats, and BHD promoted PNI recovery by regulating cells to accelerate cell cycle through p27Kip1 protein.

**Figure 2.**
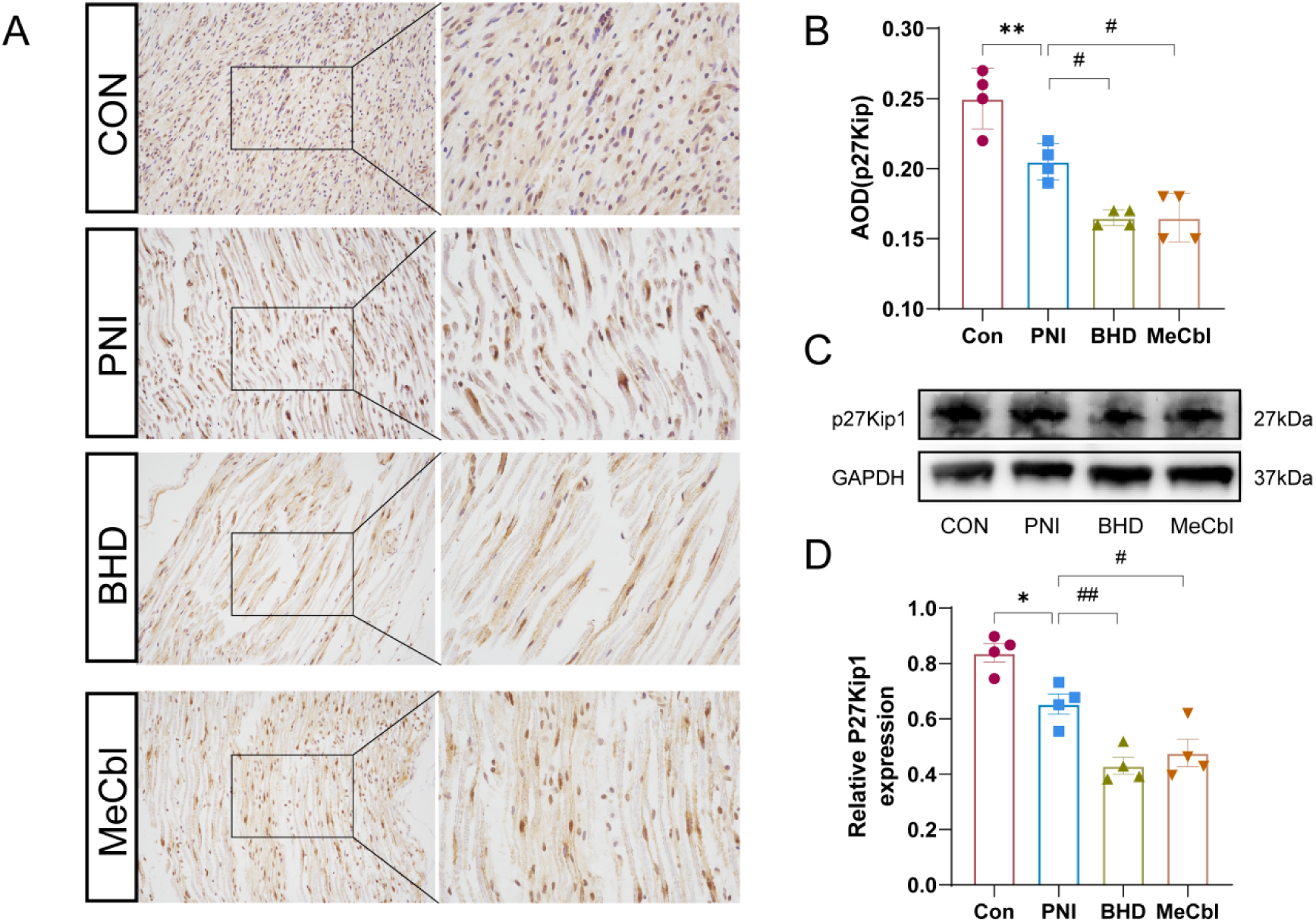
BHD decreases p27Kip1 protein expression in PNI rats. (A, B) Immunohistochemical detection of BHD inhibited p27Kip1 expression in sciatic nerve of PNI rats (n = 4). (C, D) Western blotting of BHD inhibited p27Kip1 expression in sciatic nerve of PNI rats (n = 4). All data were analyzed by one-way ANOVA, ***P< 0.001,**P< 0.01, *P < 0.05 vs. Con group, ###P < 0.01, ##P < 0.01, #P < 0.05 vs. PNI group.

### Identification of in vivo chemical constituents of BHD extracts

UPLC-Q-TOF-MS/MS was used for the preliminary identification of the main chemical constituents of BHD extract. The in vivo constituents of BHD extract were identified by the retention time, exact molecular mass, and MS/MS fragmentation pattern of reference standards in positive and negative ion modes with the literature, and a total of 40 compounds were identified (Table 1, Fig3). 10 of these compounds were from *Astragali Radix*,, 9 from *Angelica Sinensis Radix*, 9 from *Paeonia lactiflora*, 5 from *Pheretima*, and 8 from *Carthami Flos*. Kaempferol was mainly derived from *Paeonia lactiflora*, and flavonoids were mainly derived from *Astragalus membranaceus* and *Carthami Flos*.

**Table 1.**
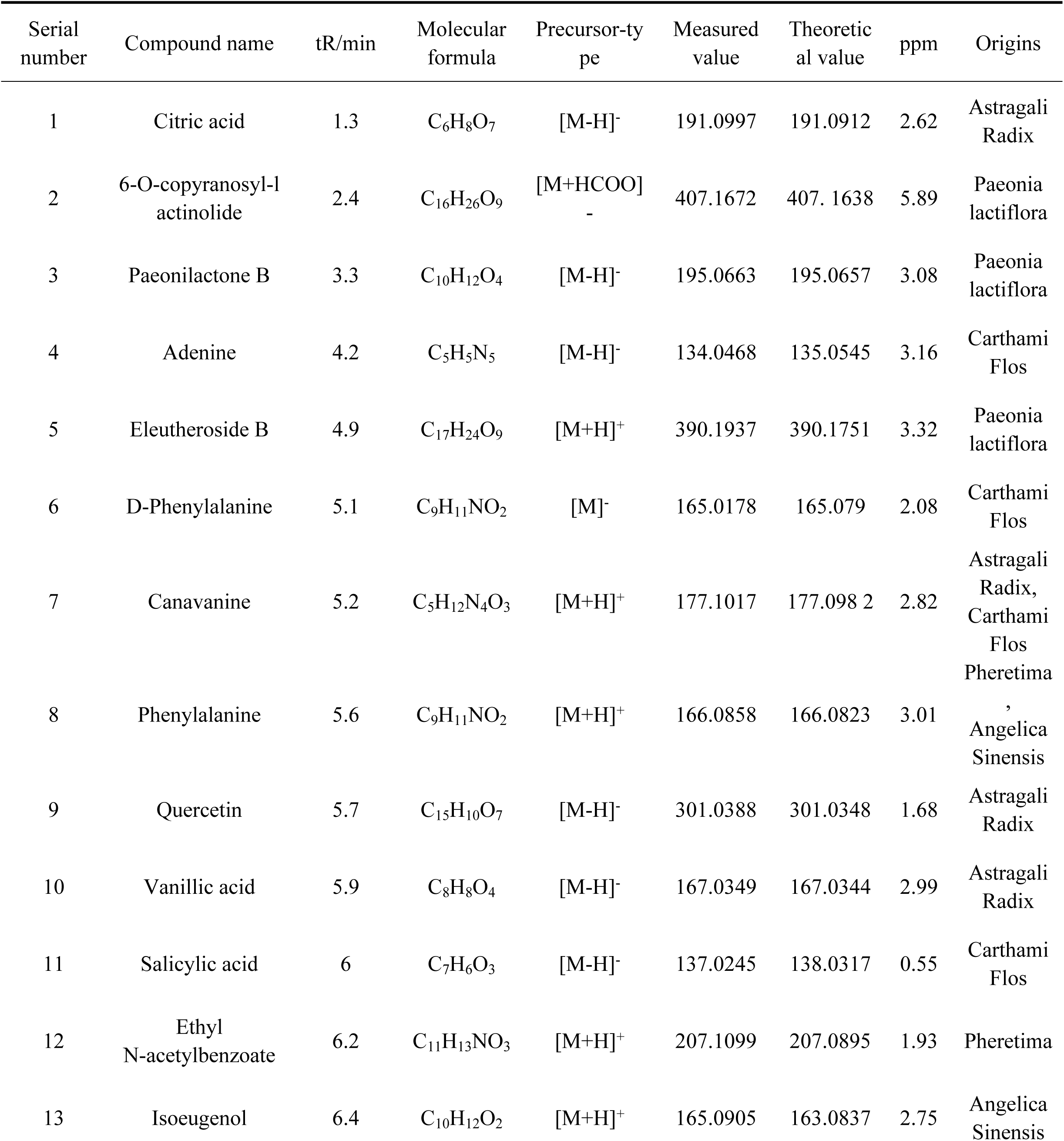
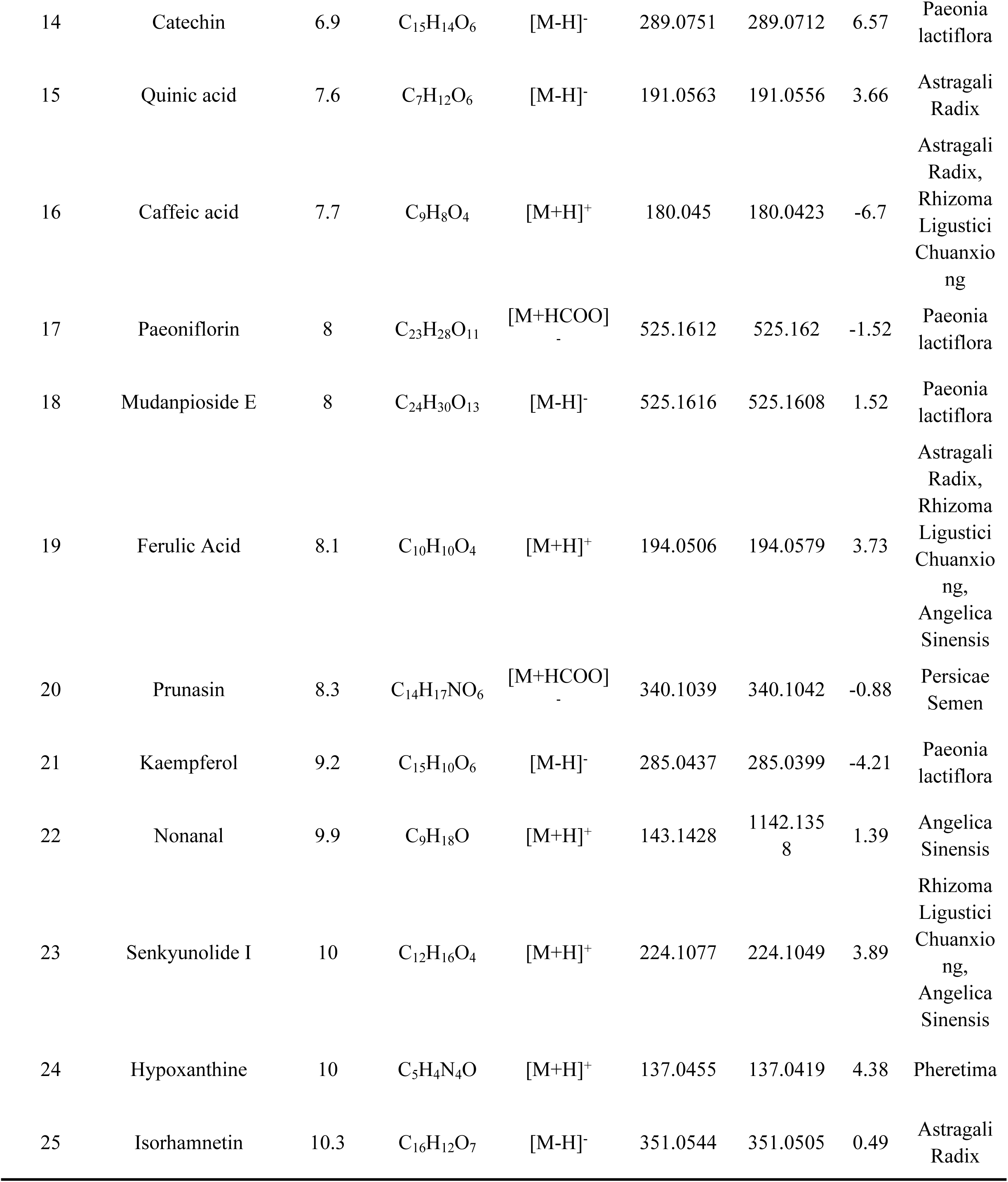
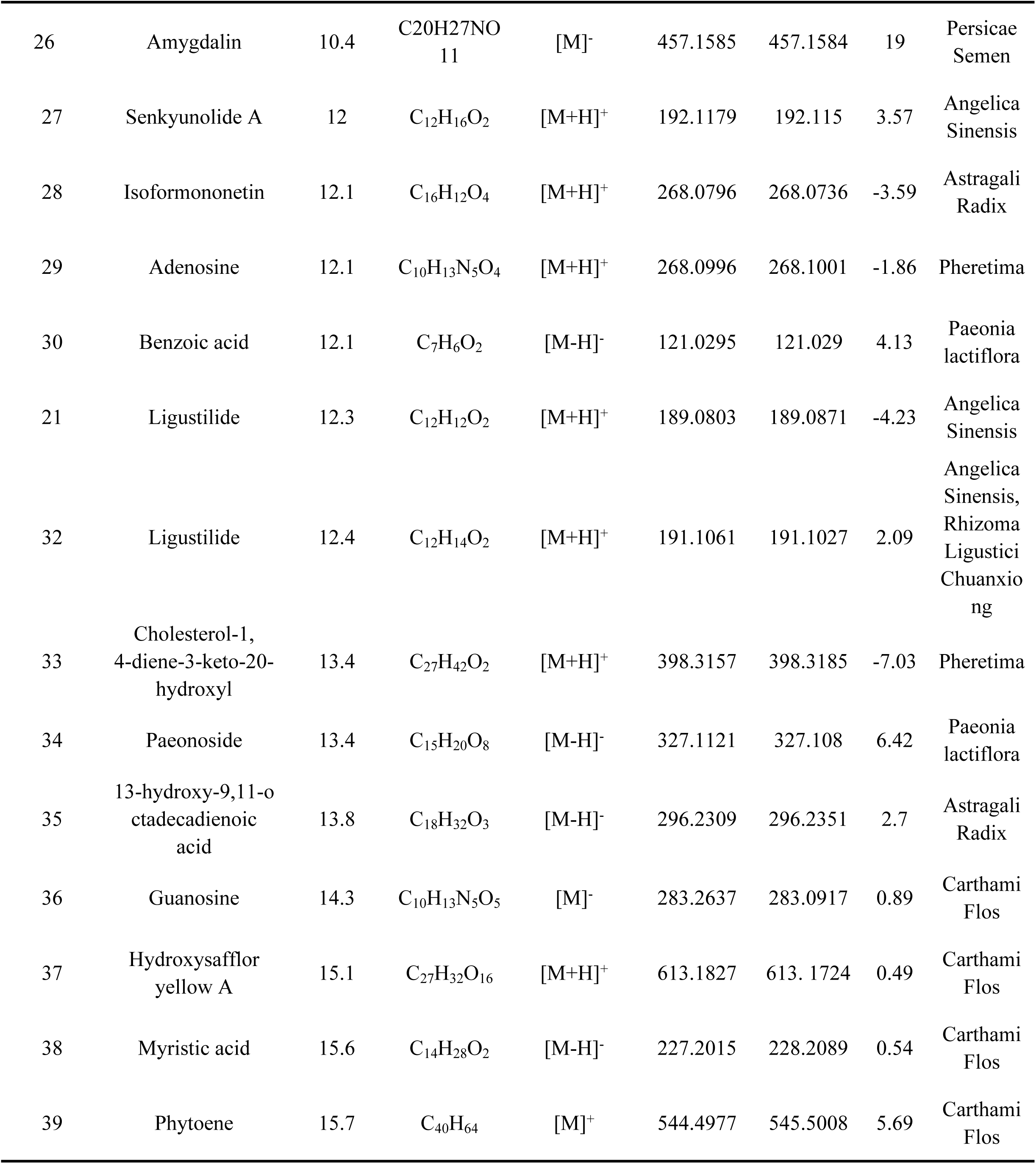
Identification of chemical constituents of BHD extract.

**Figure 3.**
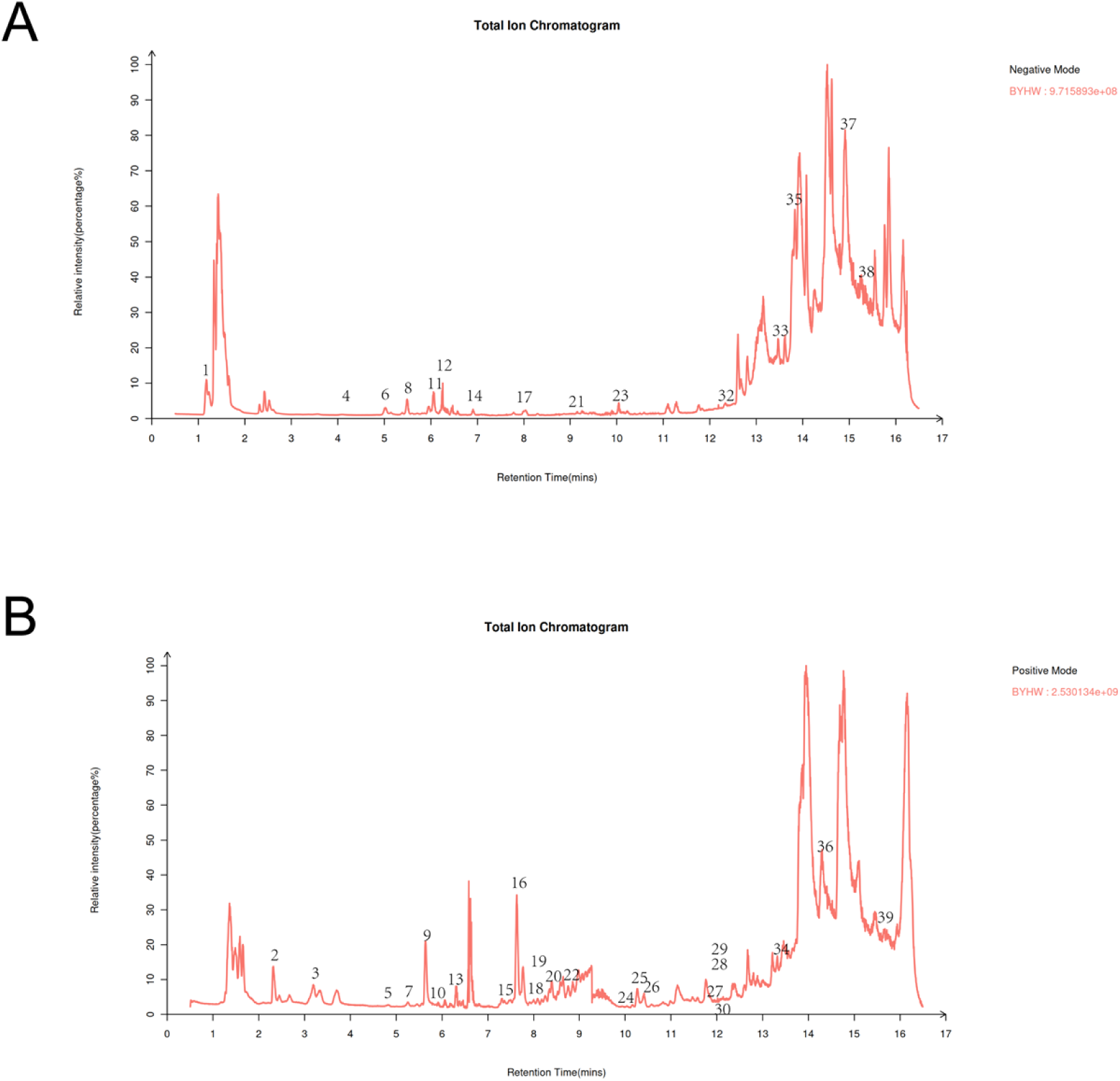
Identification of in vivo chemical constituents of BHD extracts. (A) TIC ion flow chromatogram of BHD in negative ion mode. (B)TIC ion flow chromatogram of BHD in positive ion mode.

### Pharmacologic analysis of BHD ameliorating sciatic nerve injury network in PNI rats

In the in-depth study of BHD treatment of PNI related targets, we made use of GeneCards, OMIM, DisGeNET and other database resources, and successfully identified a total of 1361 potential associated targets after screening and comparison. Further, in order to identify the associations between these targets and BHD, we performed target mapping analysis and finally identified 137 core targets that were shared by PNI and BHD. The intersection between PNI related targets and BHD targets was demonstrated by constructing a Venn diagram. (Fig. 4A). In order to further explore the potential molecular mechanism of BHD in the treatment of PNI, we introduced the selected cross targets into the STRING database to construct the protein-protein interaction network. Subsequently, PPI network maps of these targets were constructed using Cytoscape v3.9.1. The network consists of 137 nodes, which are closely connected through 2260 edges, forming a complex interactive network. In the network diagram, node color shades are designed to visually reflect the importance of each target as a core target and its probability of becoming a key regulatory factor. The results suggest that AKT1, GAPDH, ALB,1L1B, ESR1, EGFR, STAT3 and other targets are involved in the treatment of PNI by BHD (Table 2, Fig. 4A).

**Figure 4.**
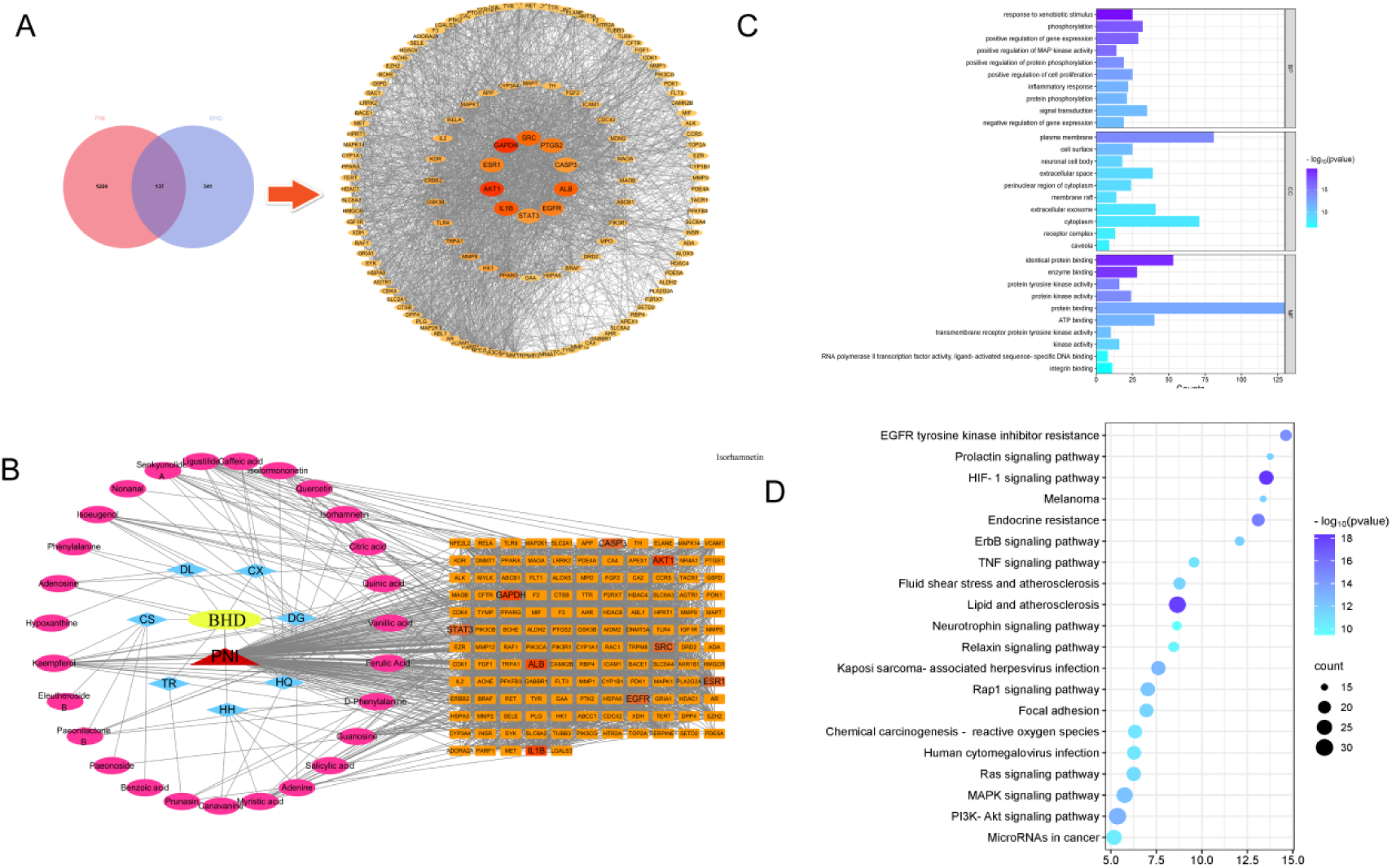
Pharmacologic analysis of BHD ameliorating sciatic nerve injury network in PNI rats. (A) BHD and PNI-related targets and protein-protein interaction network of common BHD and PNI-related targets. Node, target protein; Line, interaction between. (B) Drug-compound-disease-target network. The yellow hexagon represents BHD. The red triangle represents PNI. The blue diamond represents the formula composition of BHD. The pink circles represent active compounds contained in BHD. The orange squares represent the targets. (C) Top 10 GO terms enriched in genes. (D) Top 20 KEGG pathways enriched in genes.

**Table 2.** Top ten targets in the PPI network.

| Name | Degree | Betweenness Centrality | Closeness centrality |
| --- | --- | --- | --- |
| AKT1 | 108 | 0.01090354 | 0.82781457 |
| GAPDH | 108 | 0.04160318 | 0.84693878 |
| ALB | 95 | 0.00923318 | 0.76875 |
| IL1B | 90 | 0.03256136 | 0.77906977 |
| EGFR | 89 | 0.01459326 | 0.775 |
| STAT3 | 85 | 0.0032374 | 0.625 |
| CASP3 | 84 | 0.00973063 | 0.7254902 |
| SRC | 81 | 0.01065888 | 0.66666667 |
| ESR1 | 79 | 0.01340603 | 0.712 |
| PTGS2 | 76 | 0.01478633 | 0.8 |

In view of the fact that BHD is composed of multiple components, in this study, a comprehensive drug-compound-disease target network was constructed using Cytoscape v3.9.1 software platform to further analyze and clarify the active components of BHD that play a major role in the treatment of PNI. By analyzing this complex network, we reveal that it consists of a total of 173 unique nodes and 3,678 closely connected edges. The top ten core compounds obtained by topological analysis and sorted by degree value were Ferulic Acid, Kaempferol, Ligustilide, Guanosine, Myristic acid, Isoeugenol, Senkyunolide A, Paeonilactone B, Isorhamnetin and so on (Table 3, Fig. 4B). Therefore, these BHD compounds are considered to be effective compounds for the treatment of PNI.

**Table 3.**
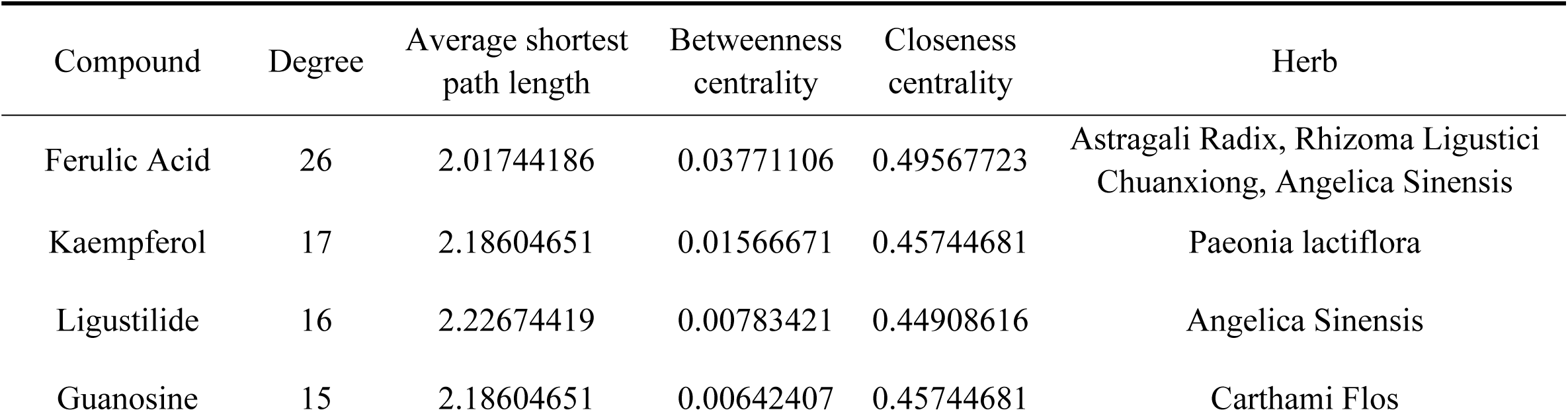
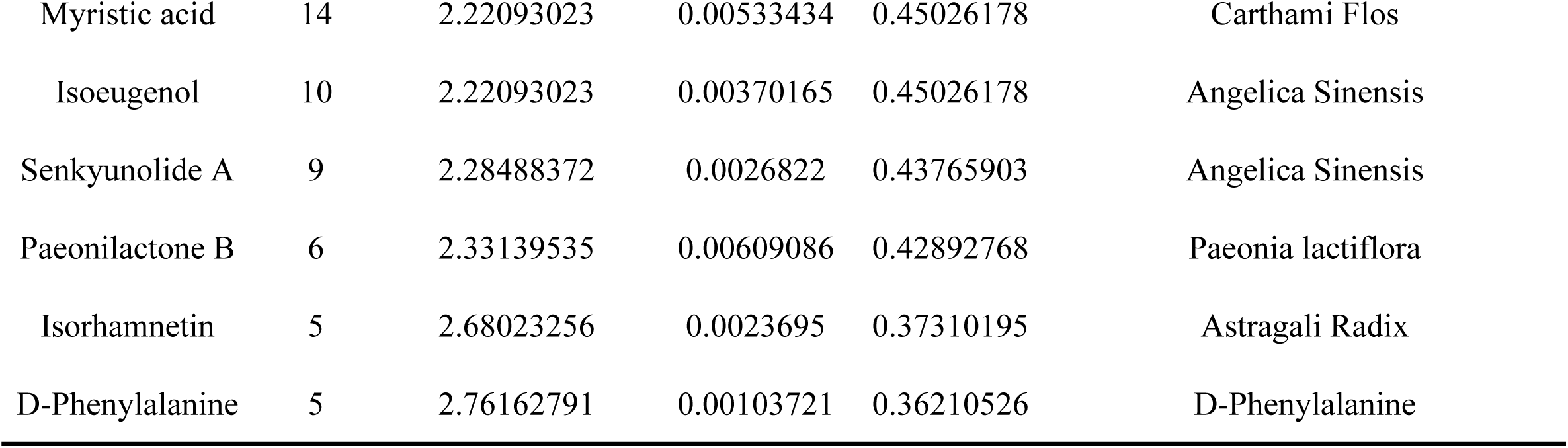
Top ten compounds in the drug-compound-disease-target network.

| Compound | Degree | Average shortest path length | Betweenness centrality | Closeness centrality | Herb |
| --- | --- | --- | --- | --- | --- |
| Ferulic Acid | 26 | 2.01744186 | 0.03771106 | 0.49567723 | Astragali Radix, Rhizoma Ligustici Chuanxiong, Angelica Sinensis |
| Kaempferol | 17 | 2.18604651 | 0.01566671 | 0.45744681 | Paeonia lactiflora |
| Ligustilide | 16 | 2.22674419 | 0.00783421 | 0.44908616 | Angelica Sinensis |
| Guanosine | 15 | 2.18604651 | 0.00642407 | 0.45744681 | Carthami Flos |
| Myristic acid | 14 | 2.22093023 | 0.00533434 | 0.45026178 | Carthami Flos |
| Isoeugenol | 10 | 2.22093023 | 0.00370165 | 0.45026178 | Angelica Sinensis |
| Senkyunolide A | 9 | 2.28488372 | 0.0026822 | 0.43765903 | Angelica Sinensis |
| Paeonilactone B | 6 | 2.33139535 | 0.00609086 | 0.42892768 | Paeonia lactiflora |
| Isorhamnetin | 5 | 2.68023256 | 0.0023695 | 0.37310195 | Astragali Radix |
| D-Phenylalanine | 5 | 2.76162791 | 0.00103721 | 0.36210526 | D-Phenylalanine |

In order to further explore the mechanism of BHD in the process of affecting PNI, we used the DAVID database to conduct detailed bioinformatics analysis on the selected core targets. The focus is specifically on the classification of Biological Process (BP), Cellular Component (CC), and Molecular Function (MF) enriched by these targets. In this analysis, 452 GO entries with statistical significance (p<0.05) were identified by consensus. P-value filtering was carried out on the top ten GO analysis entries under each category.(Fig. 4C). The results showed that in the BP category, BHD mainly affected response to xenobiotic stimulus,phosphorylation, positive regulation of gene expression andpositive regulation of cell proliferation. In addition, in the CC category, BHD is closely related to plasma membrane, cell surface, neuronal cell body, extracellular space and perinuclear region of cytoplasm. In MF category, the targets are associated with identical protein binding, enzyme binding, protein tyrosine kinase activity, protein binding and sequence-specific DNA binding. In addition, we performed KEGG pathway enrichment analysis (p<0.05) on the core targets, and a total of 152 pathways were identified. The top 20 signaling pathways were selected (Figure 4D), among which the HIF signaling pathway could affect the cell cycle by influencing p27Kip1 expression through the PI3K-AKT-mTOR-HIF-1α pathway, which might be involved in the treatment of PNI by BHD.

### BDH promotes sciatic nerve repair in PNI rats through PI3K-AKT-mTOR-HIF-1a signaling

To further verify that BHD promoted sciatic nerve repair in PNI rats through PI3K-AKT-mTOR-HIF-1a signaling, experiments were performed to detect sciatic nerve-related protein expression in rats. The results showed that P-PI3K(*P*<0.05), P-Akt (*P*<0.01), P-mTOR (*P*<0.01), and HIF-1α (*P*<0.01) protein expression was increased in PNI rats compared with the Con group (Fig. 5A-E), and P-PI3K (*P*<0.01), P-Akt (*P*<0.05), P-mTOR (*P*<0.01), and HIF-1α (*P*<0.01) protein expression was increased after BHD treatment compared with the PNI group (Fig. 5A-E). P-PI3K (*P*<0.001), P-Akt (*P*<0.001), P-mTOR (*P*<0.05) protein expression was increased after methylcobalamin treatment compared with the PNI group (Fig. 5A-E). The results demonstrated that BDH promoted sciatic nerve repair in PNI rats by regulating cell cycle progression through PI3K-AKT-mTOR-HIF signaling.

**Figure 5.**
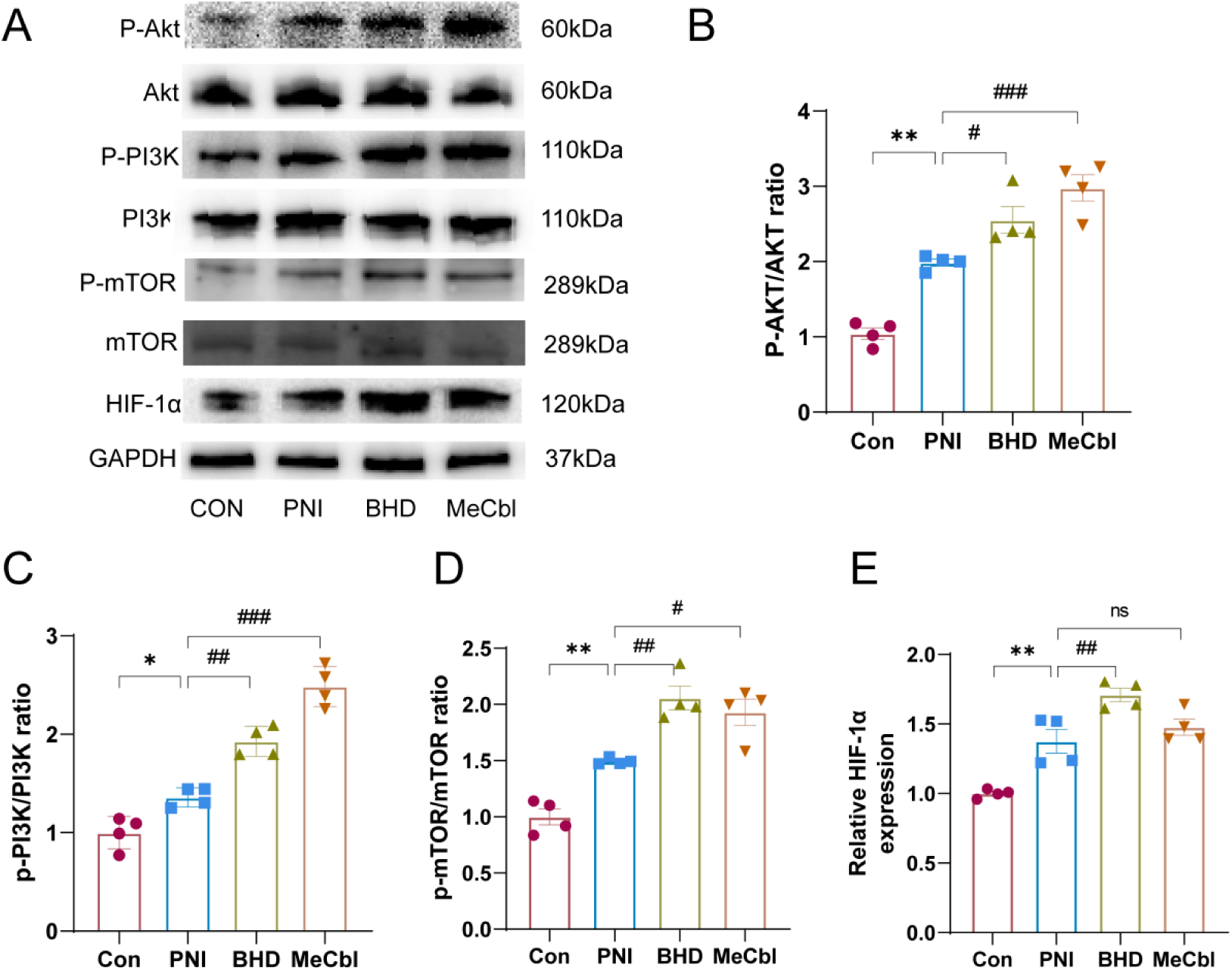
BDH promotes sciatic nerve repair in PNI rats through PI3K-AKT-mTOR-HIF-1α signaling. (A) Western blotting assays were performed to detect the expression levels of p-PI3K, PI3K, p-AKT, AKT, p-mTOR, mTOR and HIF-1α. The p-JNK/JNK. (B) Relative expression ofp-AKT. (C) Relative expression of p-PI3K. (D) Relative expression of p-mTOR. (E) Relative expression of HIF-1α. All datas were calculated by grayscale analysis, ***P< 0.001,**P< 0.01, *P < 0.05 vs. Con group, ###P < 0.01, ##P < 0.01, #P < 0.05 vs. PNI group.

## Discussion

PNI is a clinical syndrome resulting from trauma to the peripheral nerve trunk or its branches, leading to motor, sensory, and autonomic dysfunction in the corresponding areas innervated by the damaged nerve. The treatment of PNI has always been a difficult problem in today’s clinic^[26]^, although many scholars have devoted themselves to the study of promoting peripheral nerve regeneration, both medication and surgical repair usually fail to completely regenerate the cells, and even cause muscle atrophy, joint contracture, deformation^[27, 28]^. Multicomponent traditional Chinese medicine (TCM) has unique advantages in the treatment of chronic complex diseases, and TCM has thousands of years of clinical experience in the treatment of PNI.BHD was first published in the “Yilin Gaicuo” by Wang Qingren, a famous medical doctor of the Qing Dynasty, and is widely used in sciatica, polyneuritis, diabetic peripheral neuropathy, coronary heart disease, stroke and other neurological injuries^[29, 30][31–34]^and has been proved to have a good efficacy in long-term studies. The formula uses a large dose of *Astragali Radix* together with *Angelica Sinensis Radix* Tail to promote the operation of qi and blood and repair skeletal muscle damage. At the same time, *Paeonia lactiflora*, *Rhizoma Ligustici Chuanxiong*, *Persicae Semen*, *Carthami Flos*, and *Pheretima* are used to promote blood circulation. By combining qi tonifying herbs and blood activating herbs in appropriate proportion, this formula can relieve pain and promote repair of limb injury without side effects. Although BHD was first used to treat stroke^[32, 35]^, its effects of tonifying qi, invigorating blood and clearing collaterals can also have a good therapeutic effect on PNI, and its efficacy in treating PNI has been confirmed in both basic and clinical studies ^[36–38]^. In this study, we demonstrated that BHD inhibits cell cycle-associated P27KIP protein expression through the PI3K-AKT-mTOR-HIF-1a signaling pathway thereby accelerating the cell cycle to ultimately repair PNI, which not only clarifies the mechanism of action of BHD in the treatment of PNI by regulating the cell cycle, but also reveals the important clinical value of BHD in the future treatment of PNI.

In this study, we chose the clamp injury model as the model modeling method of PNI, and took Sunderland grade III injury for testing, so that the nerve function was lost, but there was some self-repair ability. Different sciatic nerve injury models correspond to different types of injuries and have different degrees of injury, among them, the clamp injury model is currently widely used in PNI experimental research as a modeling method and different clamp force and time can control the degree of nerve injury, which mainly simulates a series of changes in human peripheral nerves after suffering from compression injury^[39]^. Considering that nerve function recovery is the main purpose of nerve repair, combining the literature, we chose the inclined plate experiment and the nerve function index experiment which can directly observe the nerve function recovery of rats^[40]^, and the results showed that the nerve function of rats in the model group was significantly decreased, and the nerve function was significantly recovered after 3d, 7d, and 14d treatments with BHD. Although the results showed that the neurological function of the model group gradually recovered with the passage of time, BHD had the effect of significantly accelerating the recovery of PNI, mentioning that BHD had a better effect on promoting the recovery of neurological function in the rat PNI model. HE staining is a classic morphological test for the observation of pathomorphologic changes that occur after nerve injury and during nerve repair and regeneration. In this study, HE staining was used to observe the morphology of the sciatic nerve in rats, and the results showed that the nerve fibers of the PNI group were disorganized at the time of injury, the nerve was obviously fractured, and more tissue fragments, axon disintegration, and myelin sheath dehiscence could be seen, while the nerve fibers were arranged in a slightly more neat manner and the number of SCs was higher after BHD treatment, suggesting that BHD had a better effect on the pathologic repair after PNI.

Promoting PNI recovery through cell cycle regulation has been widely studied, Zhang et al. demonstrated that drugs can regulate cell cycle-related genes involved in cell cycle signaling pathways to promote PNI repair^[32]^, and Law et al. also demonstrated that in vivo transected sciatic nerves in rats can promote axonal regeneration through the inhibitory effect of cyclin-dependent kinases 5 (Cdk5) on Actin-Related Proteins 2/3 (Arp2/3)-dependent actin polymerization to promote axon regeneration and repair of nerve injury^[41]^. p27kip1 is a CDK regulator, which is not only an important participant in the G1/S transition, but also regulates the G2/M process and the completion of cytokinesis through cdk-dependent or cdk-independent mechanisms^[42]^. p27kip1 has been found for several years to be involved in the proliferation and differentiation of SCs after sciatic nerve extrusion to promote PNI repair through the translocation of p27 from the nucleus to the cytoplasm^[21]^. Given this, this study investigated the effect of BHD on the expression of p27kip1, a protein related to cell cycle regulation. According to previous reports we used immunohistochemistry with WB to detect p27kip1 protein expression and found that p27KIP1 was significantly decreased after BHD, although p27KTP1 expression was decreased in the model group compared with the normal group, which may be due to normal cellular repair after injury. These results suggest that BHD may promote PNI repair by regulating the cell cycle through influencing p27KTP1 expression.

In regard to the complexity of herbal components we analyzed the BHD extract in rat blood by UPLC-Q-TOF/MS technique and obtained a total of 42 components, of which kaempferol was mainly derived from Paeonia lactiflora, with anti-inflammatory and antioxidant effects^[43, 44]^, and flavonoids were mainly derived from Astragalus membranaceus and Carthami Flos, with anti-inflammatory and neuroprotective effects^[45]^. In addition, volatile oils and organic acids with antioxidant and neuroprotective effects were found in the bloodstream^[46]^. Next, by retrieving disease targets and mapping them with targets related to blood entry components and constructing PPI network, it was concluded that core targets such as AKT1, GAPDH, ALB,1L1B, STAT3 may participate in BHD’s treatment of PNI.In drug-compounddisease-target network we found that BHD compounds affect multiple targets and possess overlapping targets that may act synergistically and from the network we obtained the core compounds for BHD treatment of PNI including Prunasin, Benzoic acid, Paeonoside,Paeonilactone B, Eleutheroside B, Kaempferol, Hypoxanthine, Adenosine, Phenylalanine,Isoeugenol.To explore the mechanism of BHD for PNI, we performed GO and KEGG enrichment analysis. GO enrichment results showed that target genes were mainly enriched in: response to xenobiotic stimulus,phosphorylation, positive regulation of gene expression andpositive regulation of cell proliferation, protein binding, enzyme binding, protein tyrosine kinase activity and protein binding. KEGG enrichment analysis is mainly enriched to signaling pathways related to Pathways in cancer, HIF-1 signaling pathway, Lipid and atherosclerosis and AGE-RAGE signaling pathway in diabetic complications. Notably, HIF signaling pathway serves as a main regulator of many vital process that cell proliferation, differentiation and apoptosis^[21, 47, 48]^. HIF-1α is a transcription factor that exerts its activity under specific hypoxia^[49]^, and is widely found in mammalian cells. The signaling pathways it is involved in mainly include the PI-3K/Akt/HIF-1α pathway^[50]^, the SENP1/HIF-1α signaling pathway^[51]^, the HIF-1α/BNIP3/Bcline-1 signaling pathway^[47]^ and the MAPK/HIF-1α signaling pathway^[52]^. In addition, it has been found that increased HIF-1α protein expression increases Cyclin D1, CDK2 and p21 protein expression to promote cell proliferation^[53, 54]^. Elizabeth et al. also reported that stable expression of HIF-1α in vascular smooth muscle cells regulates vasoconstriction by limiting myosin light chain phosphorylation and regulates vascular remodeling through p27 induction in a mouse model of hypoxia^[55]^. These results emphasize the association between HIF-1α and cell cycle regulation.Finally, we tested the expression of proteins on HIF-1 Signaling Pathway of network pharmacology through WB experiment, and found that p-PI3K, p-Akt, p-mTOR, and HIF-1a levels in peripheral nerve tissue of rats after BHD treatment were significantly increased. This is preliminary evidence that BHD treats PNI by regulating the cell cycle.In conclusion, our experiments demonstrated that BHD regulates the cell cycle by activating the PI3K-AKT-mTOR-HIF-1a pathway, promoting peripheral nerve repair and ultimately improving PNI. This study provides a new perspective for the development of PNI combination drugs for clinical prevention and treatment, and also provides theoretical support for the future clinical application of PNI.

Although we demonstrated the efficacy and mechanism of BHD in the treatment of PNI, there are still limitations in this study. First of all, from the pharmacodynamic results, we obtained that the efficacy of BHDH was slightly better than that of BHDL group for follow-up experimental verification, but the dose-effect relationship of BHD in the treatment of PNI remains to be further explored. Second, some compounds whose disease target genes may not be included in the public database remain to be further refined.

### CRediT authorship contribution statement

**Honghui Li:**Writing–review& editing,Writing–original draft,Visualization, Validation,Supervision,Formal analysis,Data curation,Conceptualization,Funding acquisition.**Xudong Zhou,Pengtao Huang and Xiaoqing Zhang**:Writing–original draft,Visualization,Validation,Supervision,Methodology,Software,Formal analysis,Data curation, Conceptualization.**Yuping Gao**: Writing–original draft, Visualization, Validation, Supervision, Formal analysis, Data curation, Conceptualization.**Bifeng Fu**: Writing– review & editing, Writing– original draft, Visualization, Validation, Supervision, Software, Funding acquisition,Formal analysis, Data curation, Conceptualization.

## Ethics approval

All experiments were approved by the Ethics Committee of the First Hospital of Hunan University of Traditional Chinese Medicine, and all operations complied with the ethical requirements for laboratory animals (ZYFY20210507-11).

## Manuscript/data statement

This manuscript/data, or portions thereof, has not been published in another journal or the work has not been previously published elsewhere.

## Acknowledgments

This work was supported by the General Project of Hunan Provincial Administration of Traditional Chinese Medicine (No. B2024042) and the Joint Fund Project of Hunan University of Traditional Chinese Medicine (No. 34/2023).

## Data availability

The datasets used and analyzed during the current study are available from the corresponding author on reasonable request.

## Abbreviations

PNI: Peripheral nerve injury
BHD: Buyang Huanwu Decoction
TCMSP: Traditional Chinese Medicine Systems Pharmacology database and analysis platform
STITCH: Search Tool for Interacting Chemicals
CDK: cell cycle protein-dependent kinase
CDKI: cell cycle protein-dependent kinase inhibitor
PPI: protein-protein interaction
GO: Gene ontology
KEGG: Kyoto encyclopedia of genes and genomes
PVDF: Polyvinylidene difluoride
HE: Hematoxylin and Eosin
SCs: Schwann cells
p27Kip: p27 Kinase Inhibitory Protein
p27KTP1: p27 Kinase Inhibitory Protein 1
TNF-α: Tumor necrosis factor-α
HUVECs: Human Umbilical Vein Endothelial Cells
SFI: sciatic nerve function index
PI3K: Phosphorylated Phosphatidylinositol 3-Kinase
P-AKT: Phosphorylated AKT
P-mTOR: Phosphorylated Mechanistic Target of Rapamycin
HIF-1α: Hypoxia-Inducible Factor 1-Alpha

